# Genome-wide identification of putative disease resistance genes (R genes) in carrot (*Daucus carota* subsp. *sativus*) by homology-based gene prediction

**DOI:** 10.1101/2022.10.11.511714

**Authors:** Anastassia Boudichevskaia, Thomas Berner, Jens Keilwagen, Frank Dunemann

## Abstract

The cultivated carrot (*Daucus carota* ssp. *sativus*) is one of the most important root vegetable crops grown worldwide. Carrots are highly susceptible to several pests and diseases, and disease resistance is currently among the main breeding aims. The inheritance of resistance has been reported for a few carrot foliar diseases and root-knot nematodes, but no functionally characterized resistance gene (R gene) has yet been linked as a candidate gene to any resistance locus in carrot. Knowledge about the inventory of NLR genes (nucleotide-binding leucine-rich repeat receptors) and other R genes encoding transmembrane proteins such as receptor-like proteins (RLPs) and kinases (KIN) would be necessary to associate major QTLs (quantitative trait loci) identified by bi-parental QTL analyses or GWAS (genome-wide association analysis) with functional candidate R genes. In this study, we describe a combination of a genome-wide inventory of putative full-length carrot R genes based on a homology-based gene prediction approach called GeMoMa and subsequent classification by usage of the recent version of PRGdb 4.0 database (Calle-Garcia et al. 2022). A total of 320 putative carrot R genes were identified and bioinformatically characterized, including 72 newly identified gene models, that have not yet been annotated in the currently available carrot whole genome sequence. Based on the DRAGO 3 pipeline, totally 137 putative NLR genes were found, whereas 162 putative functional RLP and KIN genes were identified in the carrot genome. About one third of the R genes was found to be organized in clusters consisting only of NLR, RLP or KIN genes. To determine the evolutionary relationships of carrot R gene predictions, we generated a phylogenetic tree based on the alignment of all 320 R proteins. Three large clades (NLR, RLP and KIN) and a small clade (RLK) were identified, which reflect well the classification obtained after DRAGO 3 analysis. The presented carrot R gene inventory might be useful for resistance gene isolation, the development of (functional) molecular markers and resistance breeding in carrot.

## Background

The cultivated carrot (*Daucus carota* ssp. *sativus*) is one of the most important root vegetable crops grown worldwide. Carrot production can be affected by a wide range of pests and pathogens. At least five diseases of carrot are caused by bacterial pathogens, 36 by fungal and oomycete pathogens, two by phytoplasmas, and 13 by viruses (Du Toit et al. 2019). Additionally, seven genera of nematodes and several insect and mite pests can impair carrot cultivation (Du Toit et al. 2019). Among the most important foliar diseases are leaf blight caused by the fungus *Alternaria dauci*, leaf spot disease (*Cercospora carotae*), bacterial blight (*Xanthomonas hortorum* pv. *carotae*), and powdery mildew (*Erysiphe heraclei*) (Davis and Raid 2002). The most widespread soilborne root pathogens of carrot are cavity spot (caused by several species of *Pythium*), white mold (*Sclerotinia sclerotiorum*), and root-knot nematodes (various species of *Meloidogyne*) (Davis and Raid 2002). Storage diseases are mainly caused by necrotrophic fungi, such as *Botrytis cinerea* (gray mold), and *Mycocentrospora acerina* (liquorice rot disease).

Despite the numerous biotic stress factors that can impact carrot growth and storage, comparatively less is known about putative resistances and, therefore, about the genetics of resistances. The inheritance of resistance has been reported for a few carrot foliar diseases and root-knot nematodes. For instance, monogenic resistance to two foliar diseases, *Cercospora* leaf spot and powdery mildew, was reported, but resistance genes have not yet been mapped (Simon 2019). Several quantitative trait loci (QTLs) for leaf blight resistance (*Alternaria dauci*) were identified by Le Clerc et al. (2015, 2019). For resistance to the root knot nematode *Meloidogyne javanica* two loci called as *Mj-1* and *Mj-2* were mapped on carrot chromosome 8 at different positions (Ali et al. 2014, Parsons et al. 2015). With a single exception, no predicted or functionally characterized resistance gene (R gene) has been associated yet as a candidate gene with any resistance in carrot. A cluster consisting of four putative R genes spanning a region of only 50 kb co-localized in the same region of chromosome 8 as the *Mj-1* locus (Iorizzo et al. 2016). The occurrence of the first carrot whole genome sequence (Iorizzo et al. 2016) and the progress made in candidate gene identification by SNP-based genome-wide association analyses (GWAS) of carrot natural substances (Keilwagen et al. 2017, Ellison et al. 2018) implicate such approaches also for carrot disease resistance research. As a first basis, a preliminary characterization of the carrot R gene inventory in the carrot genome has yielded a large number of more than 600 putative R gene sequences (Iorizzo et al. 2016).

As plants are immobile, they have evolved two major types of disease resistance, basal defense and R gene mediated defense (Gururani et al. 2012). Basal defense, which can be a constituent of both non-host and host resistance, provides the first line of defense to the infection by a wide range of pathogens. R gene-mediated pathogen resistance is mainly based on effector molecules encoded by Avr (avirulence) genes of the pathogen, which are delivered directly into the plant cells during initial stage of infection. The effectors found in viral pathogens, bacteria, oomycetes, fungi, nematodes or insects cause a plant pathogen to elicit a resistance response in a host plant. These receptors coded by R genes are mainly intracellular, and they can specifically interact with pathogen effectors coded by the Avr genes following the gene-for-gene relationship model (Borelli et al. 2018). Plant R genes have been identified from numerous species through genetic approaches, and these loci are associated with resistance to a wide spectrum of pathogens. Although plants are confronted by such a large variety of pathogens with different modes of pathogenesis (e.g., biotrophs versus necrotrophs), most R genes belong to a limited set of proteins that are made up of a conserved set of domains, whose organization is used to define them (Dangl and Jones 2001, Hammond-Kosack and Parker 2003). Through a comprehensive review, Kourelis and van der Horn (2018) identified a number of 314 cloned functional plant R genes. Only 128 of the 314 gene products have a proposed mechanism, and the majority of these R genes encode cell surface or intracellular receptors (Kourelis and van der Horn 2018).

More than two thirds of plant disease resistance genes encode nucleotide-binding leucine-rich repeat receptors (NLRs), and most plant genomes carry a repertoire of hundreds of NLR genes (Van de Weyer et al. 2019). Based on their N-terminal structures, these R proteins can be further divided into two subclasses: TIR-NBS-LRR (TNL) that possesses a domain homologous to the Toll and interleukin-1 receptor (TIR), and non-TNL. Most non-TNL R proteins have a coiled-coil (CC) structure at the N terminal and are often called CC-NBS-LRR (CNL) R proteins (Wei et al 2018). The LRRs (Leucine rich repeats) represent the components that play an important role in recognition specificity, and these domains are present in the majority of R proteins (Jones 2001). There are a few further classes including groups that contain neither LRRs nor NBS domains but other functional domains, such as an intracellular serine-threonine kinase domain (Gururani et al. 2012). Transmembrane receptor proteins containing kinase and LRR domains, such as receptor-like proteins (RLP) and the receptor-like kinases (RLK), are also involved (Osuna-Cruz 2018). In the last years, several online omics platforms have been offered to facilitate the exploration, inventory and use of plant resistance genes. Among these platforms, the Pathogen Recognition Genes database (PRGdb) (Sanseverino et al. 2010) represents a user-friendly reference site and repository for plant geneticists interested in structure and putative function of genes involved in plant disease resistance.

In this study, we describe a combination of a genome wide inventory of predicted full-length carrot R genes based on a homology-based gene prediction approach called GeMoMa (Keilwagen et al. 2016) and usage of the PRGdb 4.0 database (Calle-Garcia et al. 2022) for classifying the predicted genes. A total of 320 putative carrot R genes were identified and bioinformatically characterized, including 72 gene models that have not yet been annotated in the current version of the carrot genome presented by Iorizzo et al. (2016).

## Methods

For detecting resistance genes in carrot, homology-based gene prediction was utilized using a wide range of resistance genes from diverse plant species. Based on publicly available sequences of 110 known plant resistance genes, we collected the information about resistance transcript isoforms from eight reference genome annotations including *Arabidopsis thaliana*, *Glycine max*, *Malus domestica*, *Populus trichocarpa*, *Solanum lycopersicum*, *Solanum tuberosum*, *Sorghum bicolor*, and *Vitis vinifera.* In total, 810 putative R gene sequences selected from the reference genomes were used for homology-based gene prediction using the software GeMoMa. GeMoMa (version 1.4.3) was run for each reference species in combination with RNA-seq evidence (Keilwagen et al. 2017, 2018). The predictions of all reference species were combined using the module GAF yielding 602 gene predictions for carrot. The Integrative Genomics Viewer (IGV) (Robinson et al. 2011) was used as a visualization tool for the interactive exploration of predicted transcripts and annotated *Daucus* genome loci (DCAR sequences, Iorizzo et al. 2016). After this first round of screening, 375 gene models were selected for a second round of analysis to examine if the gene models are indeed complete. Translated protein sequences were analyzed manually by a BLAST P search at NCBI using the non-redundant protein sequence database, and only predicted full-length sequences with putative R gene function were selected for the final R gene inventory. DRAGO 3, the tool for automatic annotation and prediction of plant resistance genes implemented in PRGdb 4.0 database was used to identify the LRR, Kinase, NBS and TIR domains from 60 HMM modules created for this purpose using HMMER v3 package (Calle-Garcia et al. 2022). DRAGO 3 is also able to detect CC and TM domains using COILS 2.2 and TMHMM 2.0c programs (Calle-Garcia et al. 2022). A phylogenetic analysis of 320 deduced *Daucus carota* R proteins was performed with MEGA-X after multiple sequence alignment by MUSCLE. Neighbor-joining method and n=1000 replicates for bootstrapping were applied. The software *MapChart* 2.2. (Plant Research International, Biometris, Wageningen, Netherlands) was used for R gene map visualization. R genes were considered clustered if at least three genes from the classes NLR, RLP or KIN were located within a genomic region < 200 kb.

## Results and Discussion

The objective of this research was to conduct a genome-wide (re)-identification of R genes in the cultivated carrot. Initially, we predicted 602 putative R gene models after GeMoMa analysis. After manual evaluation using the IGV browser and elimination of too short and obviously fragmentary sequences, followed by BLAST P-based analyses for full-length CDS and predicted functions, 320 putative R gene models remained (Suppl. Table 1). Among these predicted genes, 72 of them (23%) have been newly identified since they have not yet been annotated in the current version of the carrot genome (Iorizzo et al. 2016). In addition, some new R gene clusters, such as the six-gene cluster at the beginning of chromosome Chr_3 (*DcRG_082* - *DcRG_87*) or the four-gene cluster on Chr_7 (*DcRG_233* - *DcRG_236*) were detected. Based on the DRAGO 3 pipeline, 137 putative NLR genes (43%) were found, whereas 172 putative genes encoding transmembrane proteins (54%) were identified in the carrot genome (Table 1). Among the NLR genes, the largest sub-class is the NBS-LRR (NL) type, followed by the TIR-NBS-LRR (TNL) type (Table 1). A few genes showing rare or unknown domain combinations such as CK or CL (Osuna-Cruz et al. 2018) were indeed genes possibly involved in plant resistance. For instance, genes *DcRG_169* and *DcRG_192* (CTNL class) were predicted in the *D. carota* genome as TMV resistance protein N-like isoforms X1 and X2 (Table 2). Only two genes (*DcRG_079* and *DcRG_222*) were not classified by DRAGO domain recognition but were predicted by BLAST as putative EDR2s (Table 2). These carrot genes are similar (amino acid identity ~ 80%) to the *A. thaliana EDR2* resistance gene (AT4G19040, Vorwerk et al. 2007). Because loss-of-function mutations in the *EDR2* gene confer enhanced disease resistance to powdery mildew in *A. thaliana, EDR2* probably functions as a negative regulator of powdery mildew resistance (Vorwerk et al. 2007). In the work of Iorizzo et al. (2016), MATRIX-R pipeline was used to automatically retrieve, annotate, and classify carrot R genes. Based on this pipeline, 634 putative R genes were predicted, which is comparable to the 602 gene models detected after GeMoMa analysis. However, based on sequence analyses of a random sample, we noticed that not all predicted sequences appeared to represent putative functional R genes (data not shown). In the study of Iorizzo et al. (2016), 295 R genes were assigned to the NLR classes of cytoplasmic proteins, and 339 genes were classified as transmembrane receptors. The NLR sub-class with the highest number of genes was the NL class (63 genes), which is in accordance with our finding of 64 NL genes in this sub-class (Table 1). The proportion of the NLRs at the total number of R genes was 46%, which is similar to the value found after GeMoMa (43%).

**Table 1.** Predicted putative R genes identified in the carrot genome

| R protein type | Class | No. of genes |
| --- | --- | --- |
| ***NLR*** |  |  |
| CC-NBS-LRR | CNL | 19 |
| TIR-NBS-LRR | TNL | 40 |
| NBS-LRR | NL | 64 |
| CC-NBS | CN | 1 |
| TIR-NBS | TN | 10 |
| NBS | N | 3 |
| ***Transmembrane protein*** |  |  |
| Receptor-like kinase | RLK | 10 |
| Receptor-like protein | RLP | 82 |
| Kinase | KIN | 80 |
| ***Unknown domain combination***<sup>2)</sup> | CK, CL, CTNL | 9 |
| ***Other***<sup>2)</sup> |  |  |
| Enhanced disease resistance 2 | EDR2 | 2 |
|  |  | **320** |

<sup>1)</sup> according to PRGdb 4.0 database (Calle-Garcia et al. 2022)
<sup>2)</sup> for details, see Table 2

**Table 2.**
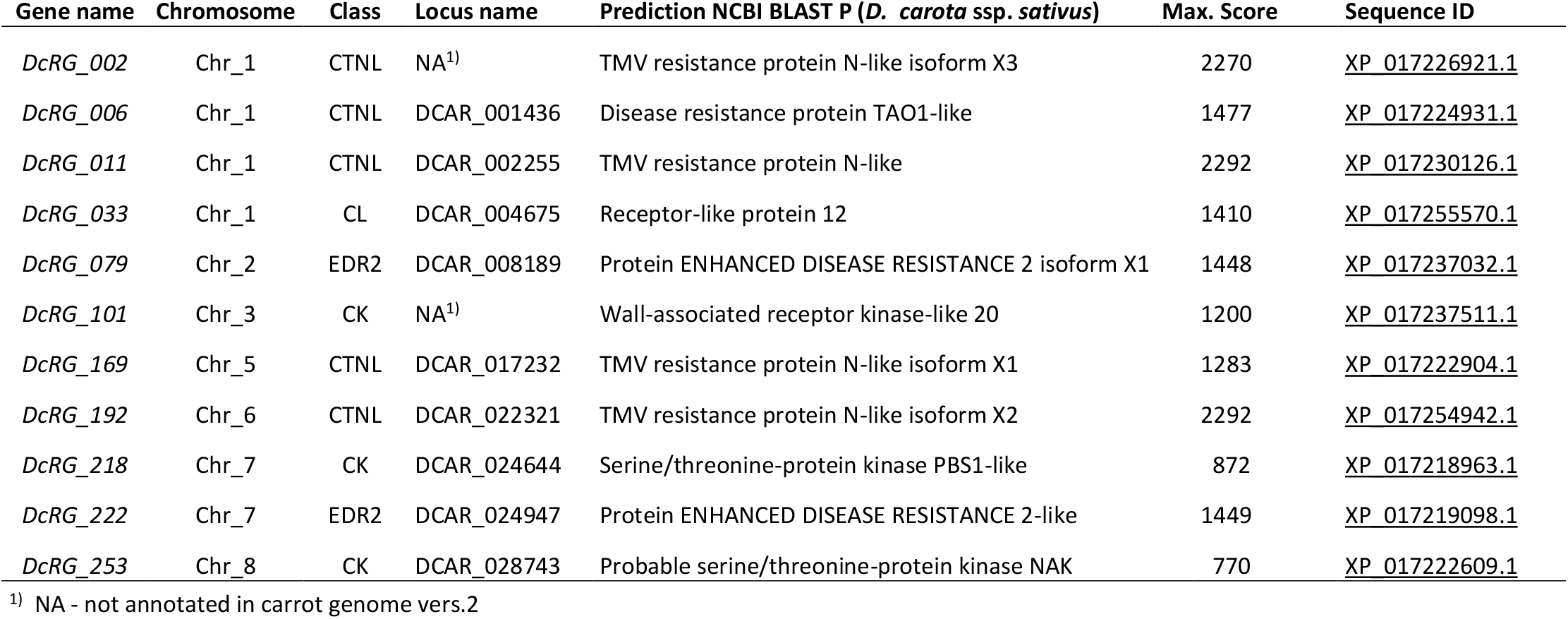
R genes with unknown domain combinations (CK, CL, CTNL) or no domain recognition (EDR2) by DRAGO 3 predicted with highest max. score by NCBI BLAST P (best hit)

**Table 3.**
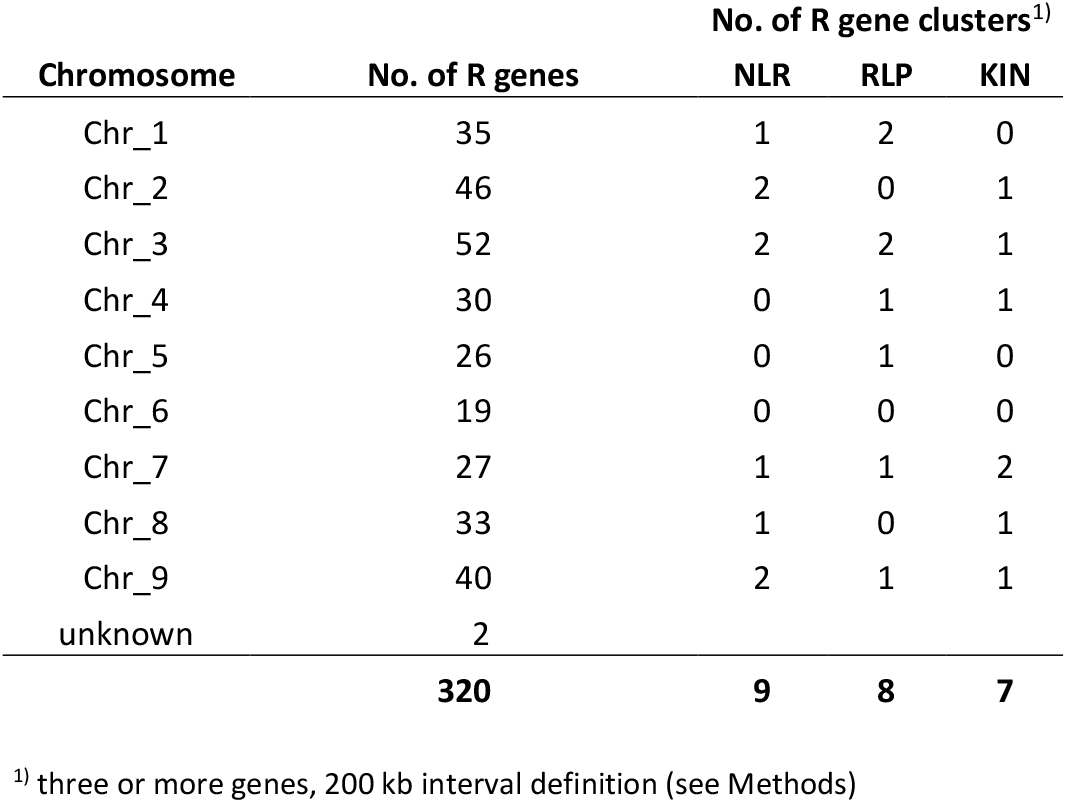
Number of predicted R genes on carrot chromosomes and number of R gene clusters according to the inventory list (Suppl. Table 1)

| Chromosome | No. of R genes | No. of R gene clusters<sup>1)</sup> |  |  |
| --- | --- | --- | --- | --- |
|  |  | NLR | RLP | KIN |
| Chr_1 | 35 | 1 | 2 | 0 |
| Chr_2 | 46 | 2 | 0 | 1 |
| Chr_3 | 52 | 2 | 2 | 1 |
| Chr_4 | 30 | 0 | 1 | 1 |
| Chr_5 | 26 | 0 | 1 | 0 |
| Chr_6 | 19 | 0 | 0 | 0 |
| Chr_7 | 27 | 1 | 1 | 2 |
| Chr_8 | 33 | 1 | 0 | 1 |
| Chr_9 | 40 | 2 | 1 | 1 |
| unknown | 2 |  |  |  |
|  | **320** | **9** | **8** | **7** |

<sup>1)</sup> three or more genes, 200 kb interval definition (see Methods)

Over the past two decades, sequencing technologies have been rapidly developed and used to assess plant-microbe interactions (Lee et al. 2015). Their use enables genome-wide analyses of NLR genes based on the NB-ARC domain (Meyers et al. 2005). Because of their importance in ecology and breeding, there has been much interest in defining inventories of NLR genes at different taxonomic levels. These efforts have revealed that the number of NLR genes across species varies from less than a hundred to over a thousand (van de Weyer et al. 2019). The number of NLR genes in flowering plants is largely variable without any clear correlation to the phylogeny, suggesting species-specific mechanisms in NLR genes expansion and/or contraction (Jacob et al. 2013). *Arabidopsis thaliana* has 159 NLR genes including 43 CNLs and 83 TNLs (Guo et al. 2011). Solanaceae plants carry more than twice the number of NLR genes than *Arabidopsis* and possess more CNLs than TNLs (Lee et al. 2015). As reviewed by Jacob et al. (2013), species with large numbers of NLRs are rice and grape with each about 460 NLR genes. Nearly 1000 NLR genes have been identified in apple, although its genome size is only approximately 740 Mb (Velasco et al. 2010). Furthermore, there is also variation in the gene copy numbers of the two subclasses, TIR-type NLRs (TNL) and non-TIR-type NLRs (CNL). In our study, we observed a CNL-type NLR to TNL-type NLR ratio of 1:2. The same ratio was found in Brassicaceae species, in potato and grapevine it was 4:1, whereas a ratio of 1:1 has been reported for apple (Borelli et al. 2018). The importance of the NLR-type R genes for plant disease resistance is also documented by the number of cloned R genes. Out of the more than 300 cloned genes, 61% encode NLRs, but only 19% of the cloned R genes encode RLPs and RLKs (Kourelis and van der Horn 2018).

The predicted carrot R genes are not evenly distributed over the nine chromosomes. The number of R genes varied between only 19 on chromosome Chr_6 and 52 on Chr_3 (Table 3). As demonstrated by the R gene map presented in Figure 1, some genomic regions appear to have no or a fewer number of R genes. Especially dense R gene regions were observed on chromosomes Chr_1 (lower end), Chr_2 (whole chromosome), Chr_3 (upper and lower part), Chr_4 (middle part), Chr_5 (lower end), Ch_7 (lower half), Chr_8 (lower half), and Chr_9 (whole chromosome). NLRs, RLPs and kinases (KIN) showed a clear tendency for clustering (Figure 1, Suppl. Table 1). The size of clusters is rather variable. The largest cluster on Chr_8 contains 11 NLRs within an interval of about 250 kb, and the second-largest NLR cluster was found on Chr_7 with 9 genes within a 340 kb-interval. Both clusters are heterogenous and contain a mixture of diverse NLRs, i.e. NL, CNL, TN, and TNL. Smaller homogenous NLR clusters were also found, as for instance the TNL-gene cluster on Chr_2 or the NL-cluster on Chr_3 (Suppl. Table 1). The in total 8 RLP and 7 KIN clusters (Table 3) were, with two exceptions on Chr_1 and Chr_9, homogenously in terms that they do not contain genes from the cytoplasmic NLR sub-classes. The three largest RLP clusters with each seven genes are located on chromosomes Chr_1, Chr_3 and Chr_5. NLR genes are known to be unevenly located in plant genomes and are often found in multigene clusters (Meyers et al. 2003). This clustered distribution likely arose by tandem duplications and further sequence divergence and depicts a large genetic reservoir for evolution of new specificities to pathogens (Duplessis et al. 2009). The homogeneous cluster type is most probably generated by tandem duplication, whereas the heterogenous cluster type is derived from ectopic duplications, transpositions, and/or large-scale segmental duplications with subsequent local rearrangements (Jacob et al. 2013). In the carrot genome, also other major R gene classes such as RLP and KIN were found to be organized in gene clusters.

**Figure 1.**
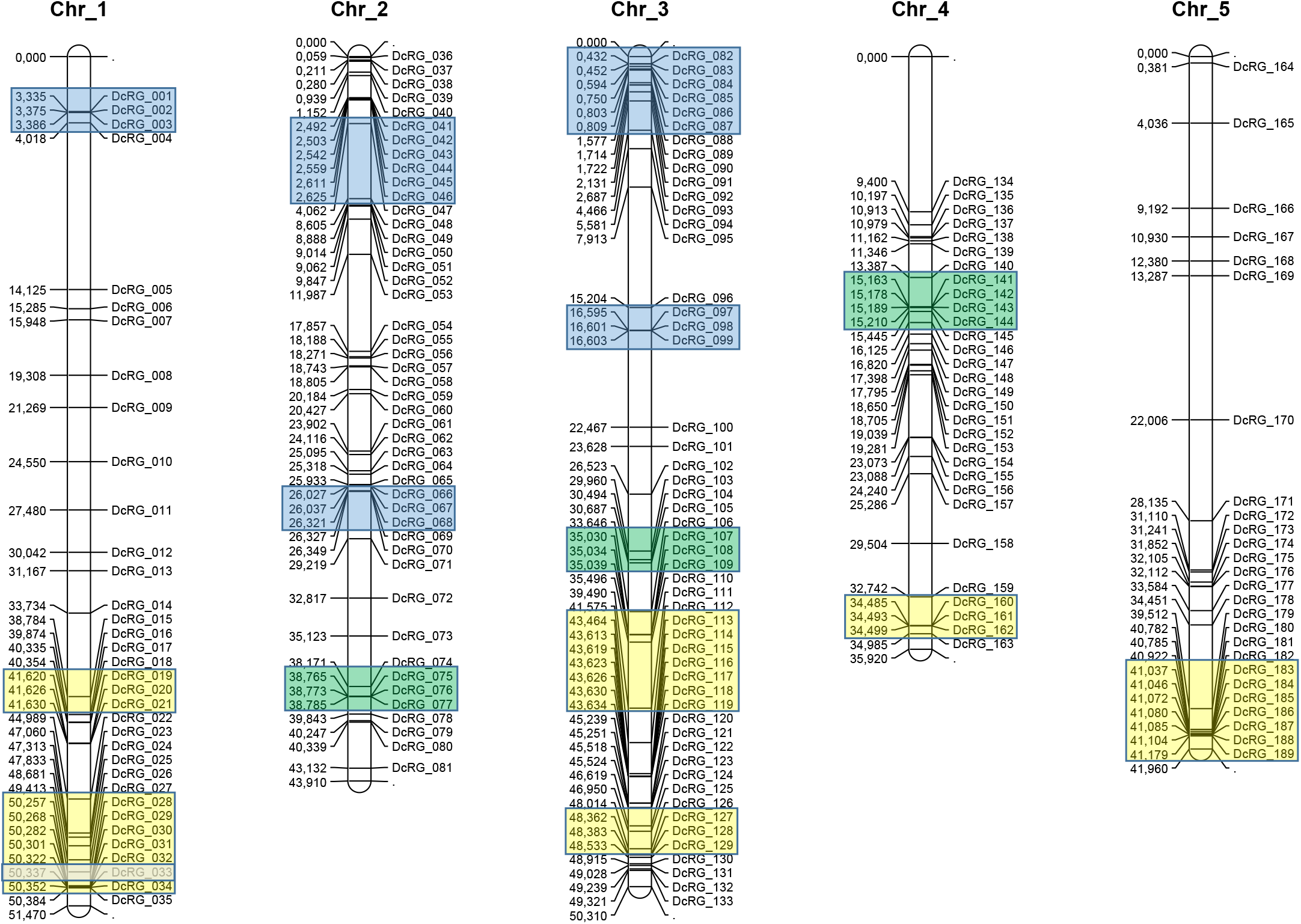
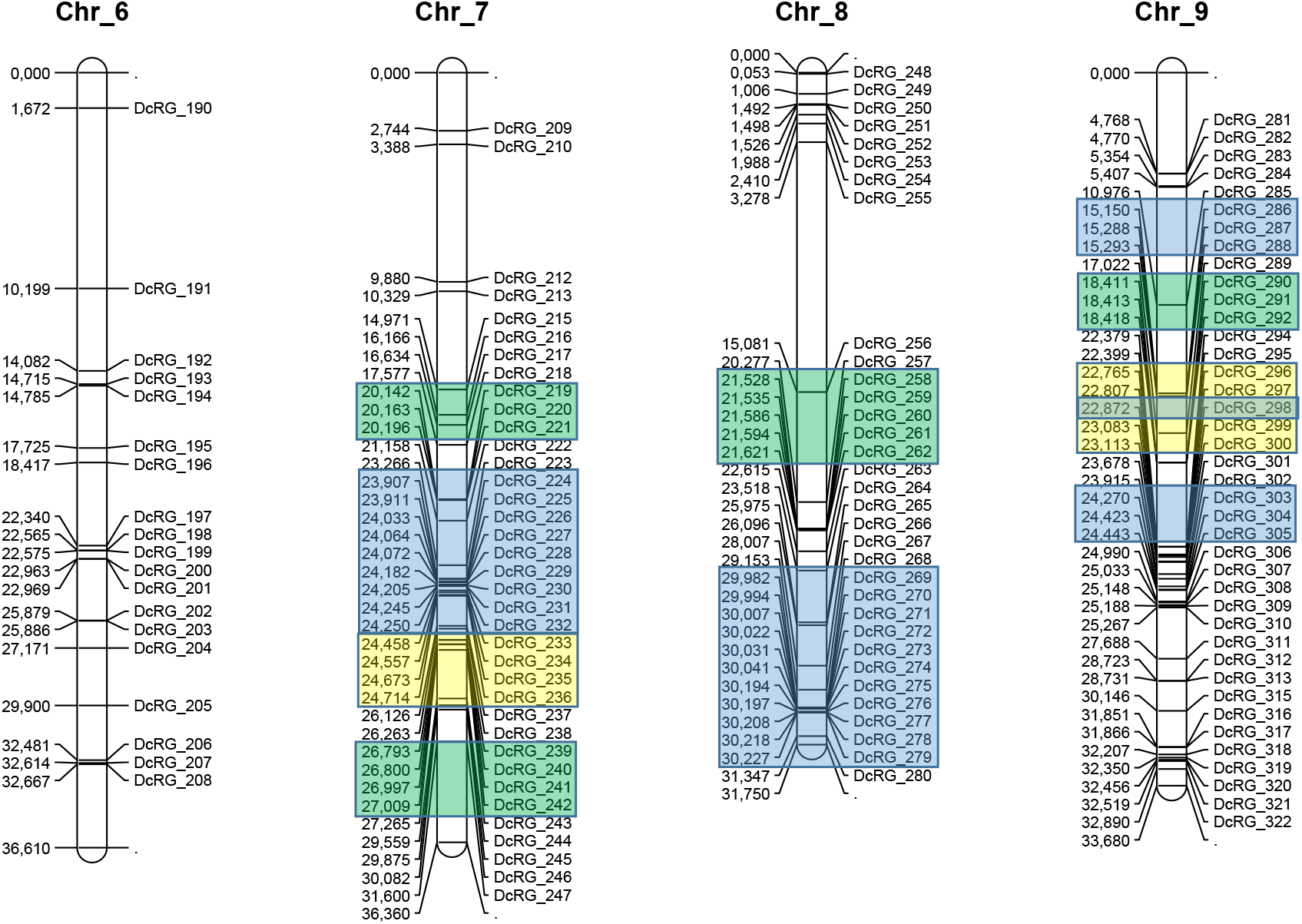
Carrot R gene map. Schematic map presentation of the genomic localization of the 320 carrot R genes listed in Suppl. Table 1. Figures on the left side of the bars show the start position of the CDS of each R gene in Mb (mega base pairs). R gene clusters are presented in coloured boxes (NLR - blue, RLP - yellow, KIN - green).

To determine the evolutionary relationships of carrot R gene predictions, we generated a phylogenetic tree based on the alignment of all 320 deduced R proteins. Three large clades (NLR, RLP and KIN) and a small clade (RLK) were identified (Fig. 2). The predicted classification of the R genes is well represented by the phylogenetic relationships. Only a single predicted NL gene (*DcRG_010*) was placed among the RLP genes. The three genes with the unknown domain combination CK (*DcRG_101*, *DcRG_218* and DcRG_253) were assigned to the KIN clade, and the single CL gene was assigned to the RLP clade, respectively. The two EDR2 genes are closely related to the two CNL genes *DcRG_093* and *DcRG_158* and function in this phylogenetic analysis like an outgroup (Fig. 2). The similarity of the sequences of clustered R genes is often reflected by their positions in the phylogenetic tree (Fig. 2). Examples are the NLR cluster on Chr_2, which contains the six NL genes *DcRG_041* - *DcRG_046*, and the eleven-gene cluster on Chr_8 with the six genes *DcRG_269* - *DcRG_274*, which also formed a common NL sub-group. The other five genes from this cluster, however, were grouped in a common TNL gene sub-group at a different position in the phylogenetic tree (Fig. 2). There are also some RLP and KIN gene clusters that obviously contain structurally similar R genes. For instance, RLP genes *DcRG_113* - *DcRG_119* (Chr_3) and KIN genes *DcRG_258 - DcRG_262* (Chr_8) were located each in the same sub-group after phylogenetic analysis indicating common evolutionary origins.

**Figure 2.**
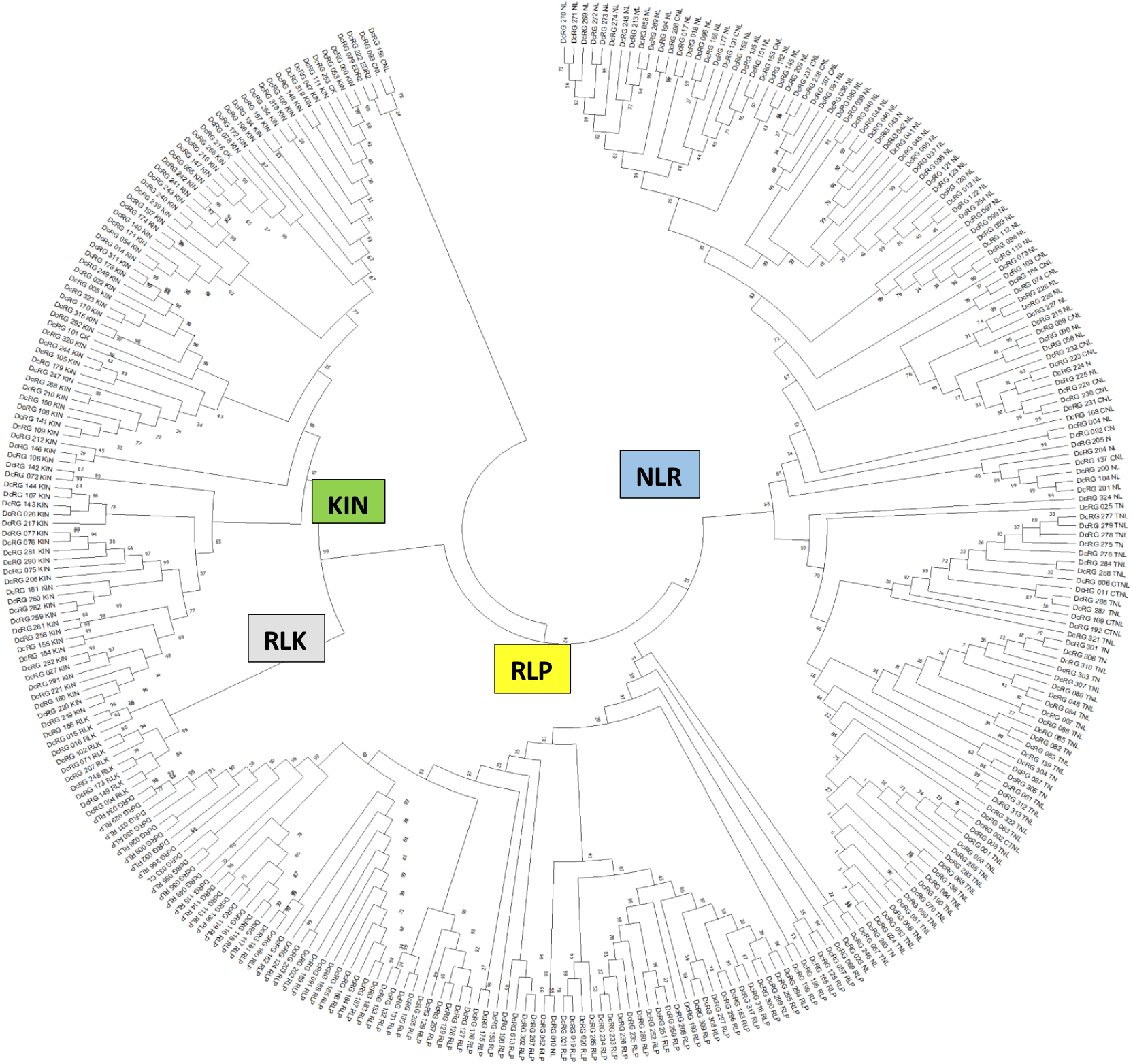
Phylogenetic analysis of 320 deduced *Daucus carota* R proteins and their grouping in clades (NLR, RLP, RLK and KIN). Multiple sequence alignment by MUSCLE (MEGA-X). Neighbor-joining method and n=1000 replicates for bootstrapping were applied (MEGA-X). Bootstrap values are shown next to each node.

## Conclusion

The presented carrot R gene inventory consisting of more than 300 putative functional genes might be useful for resistance research and resistance breeding in carrot. Studies on the genotypic and phenotypic variability of resistances against several carrot diseases, supplemented by bi-parental QTL analyses or GWAS based on larger carrot genotype collections, may identify major QTLs and associated candidate R genes. In case of conventional breeding for resistance, introgression of resistance genes from resistant wild carrot relatives by repeated backcrossing is a long-lasting process. The use of molecular markers developed directly from R genes or R gene clusters might strongly support the backcrossing program. In addition, a deeper knowledge about the structure and function of isolated R gene candidates might be used in future for biotechnological approaches, such as CRISPR/Cas9-based genome editing, with the aim to generate new or to integrate known resistance alleles in the genome of the cultivated carrot.

## Author’s contributions

AB, TB, and JK performed the bioinformatic analyses. FD performed the evaluation and classification of predictions and compiled the carrot R gene inventory. FD drafted the manuscript. AB, JK, and FD contributed to the discussion and interpretation of results and read and approved the final manuscript.

---

**Suppl. Table 1.**
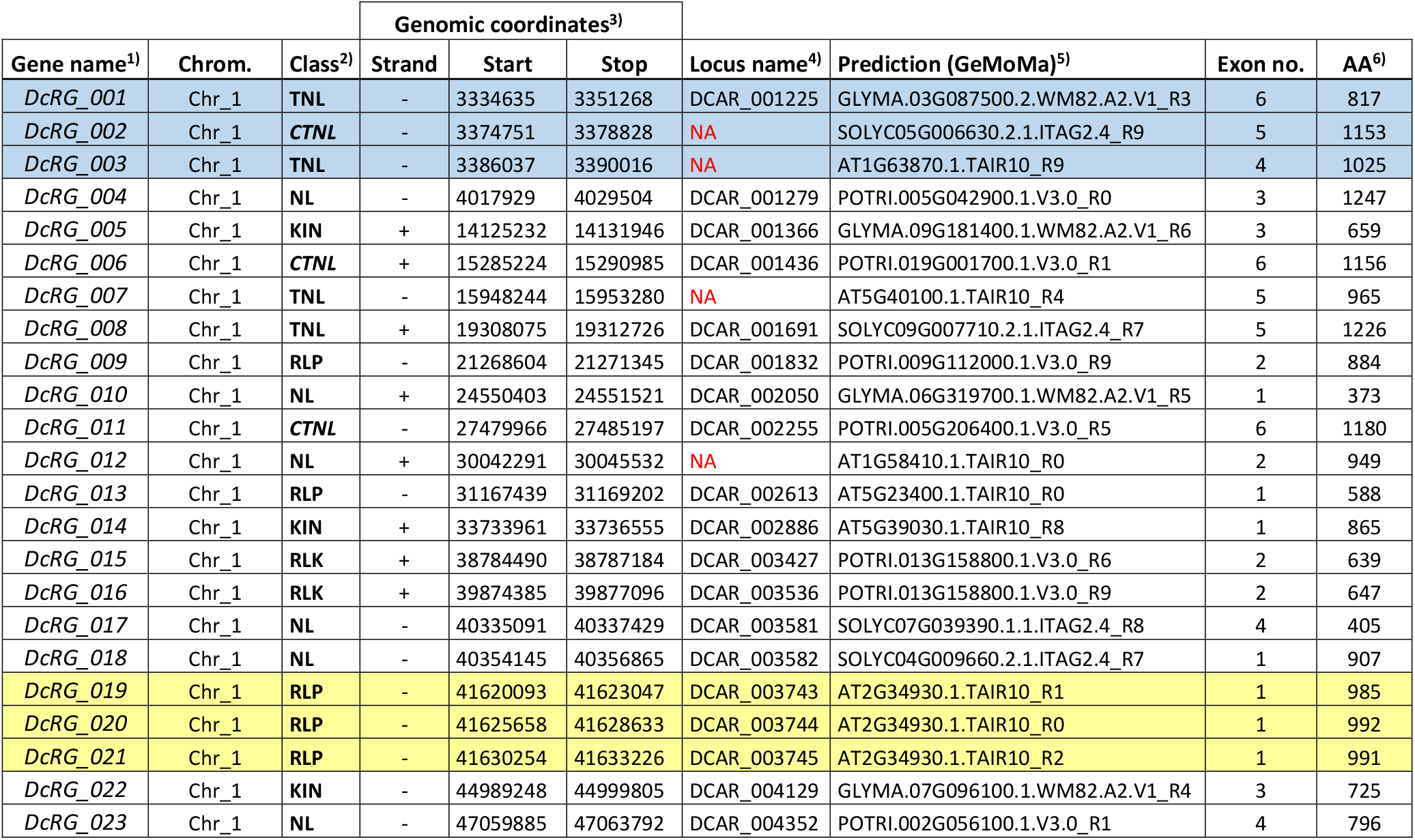
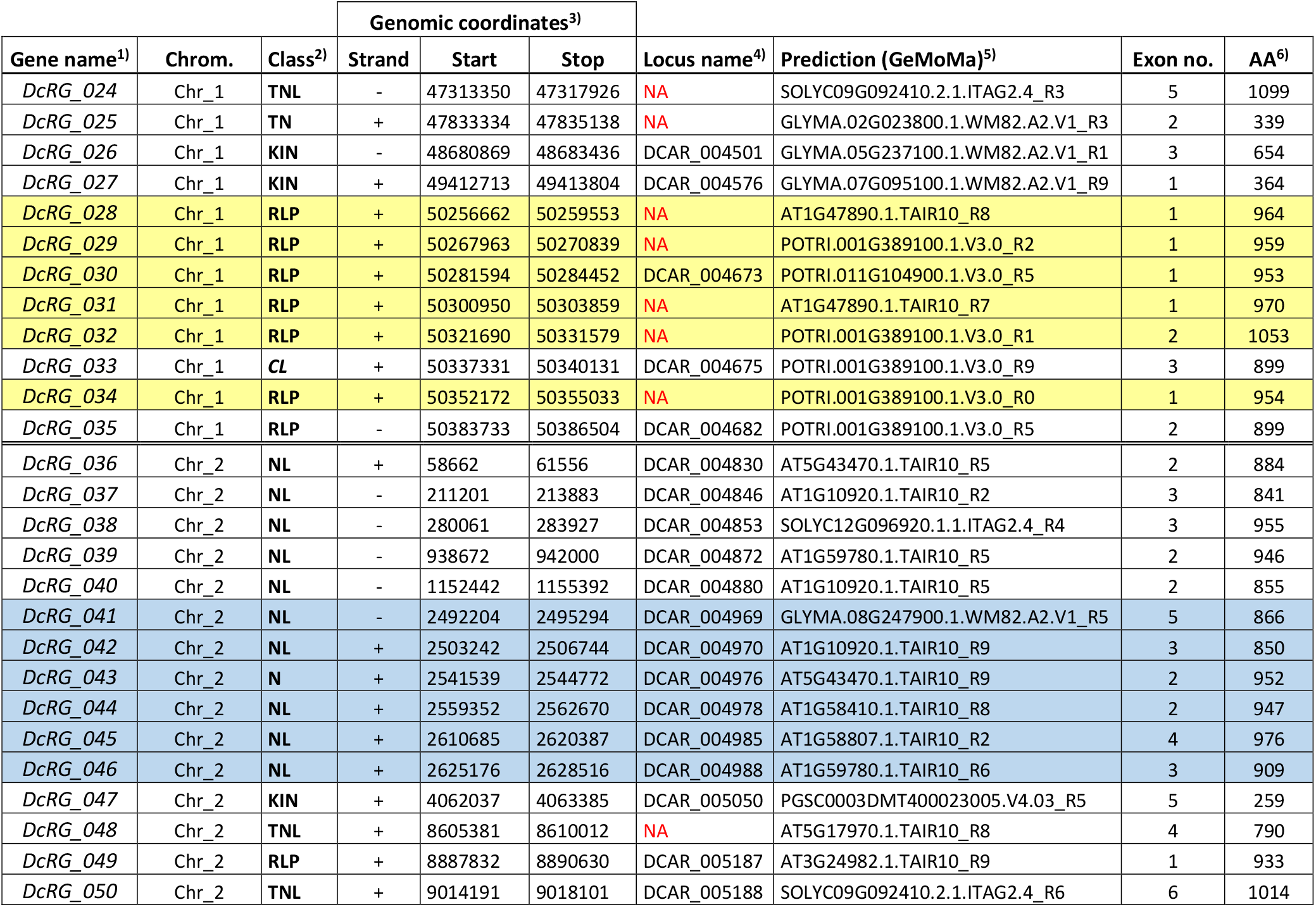
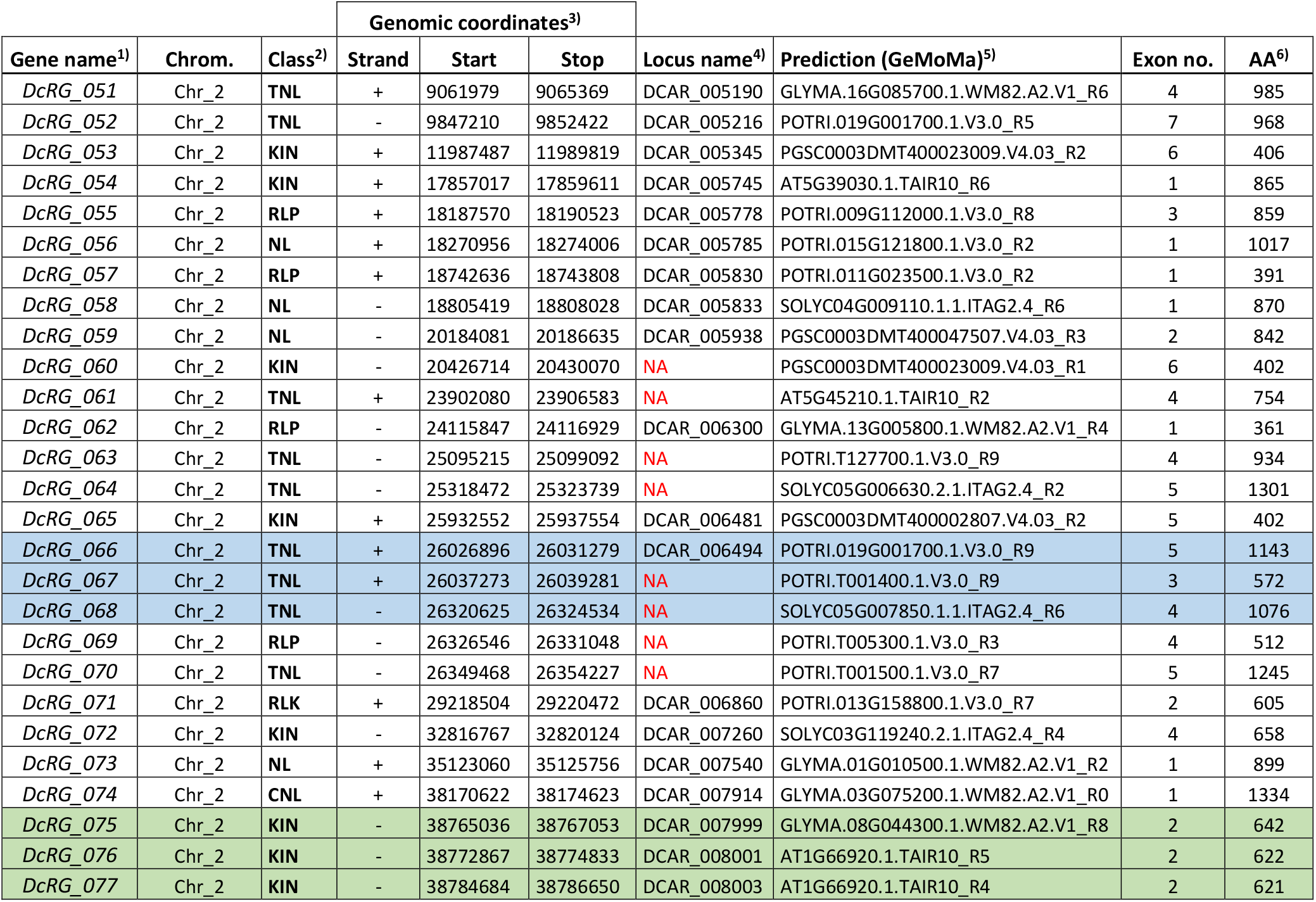
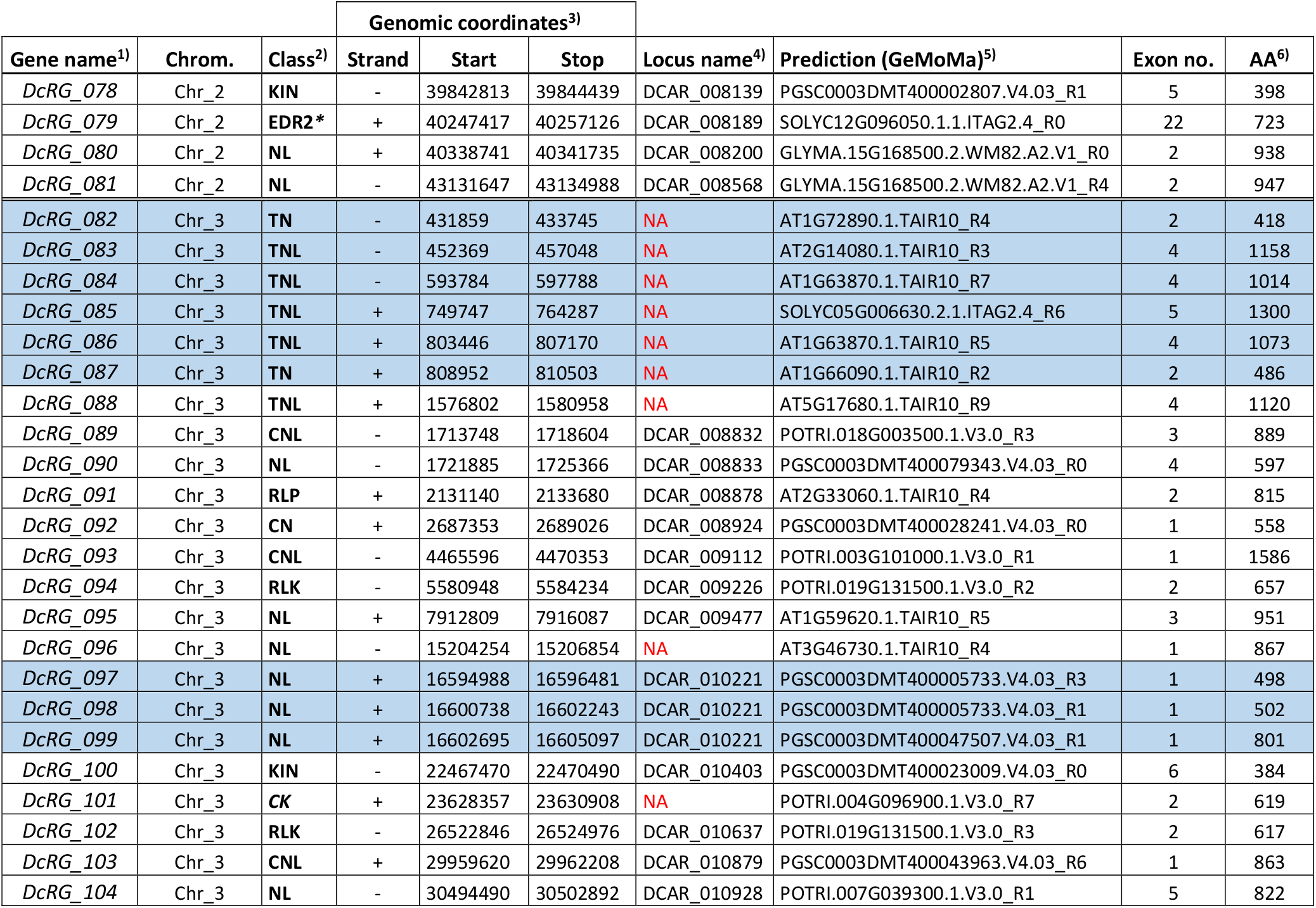
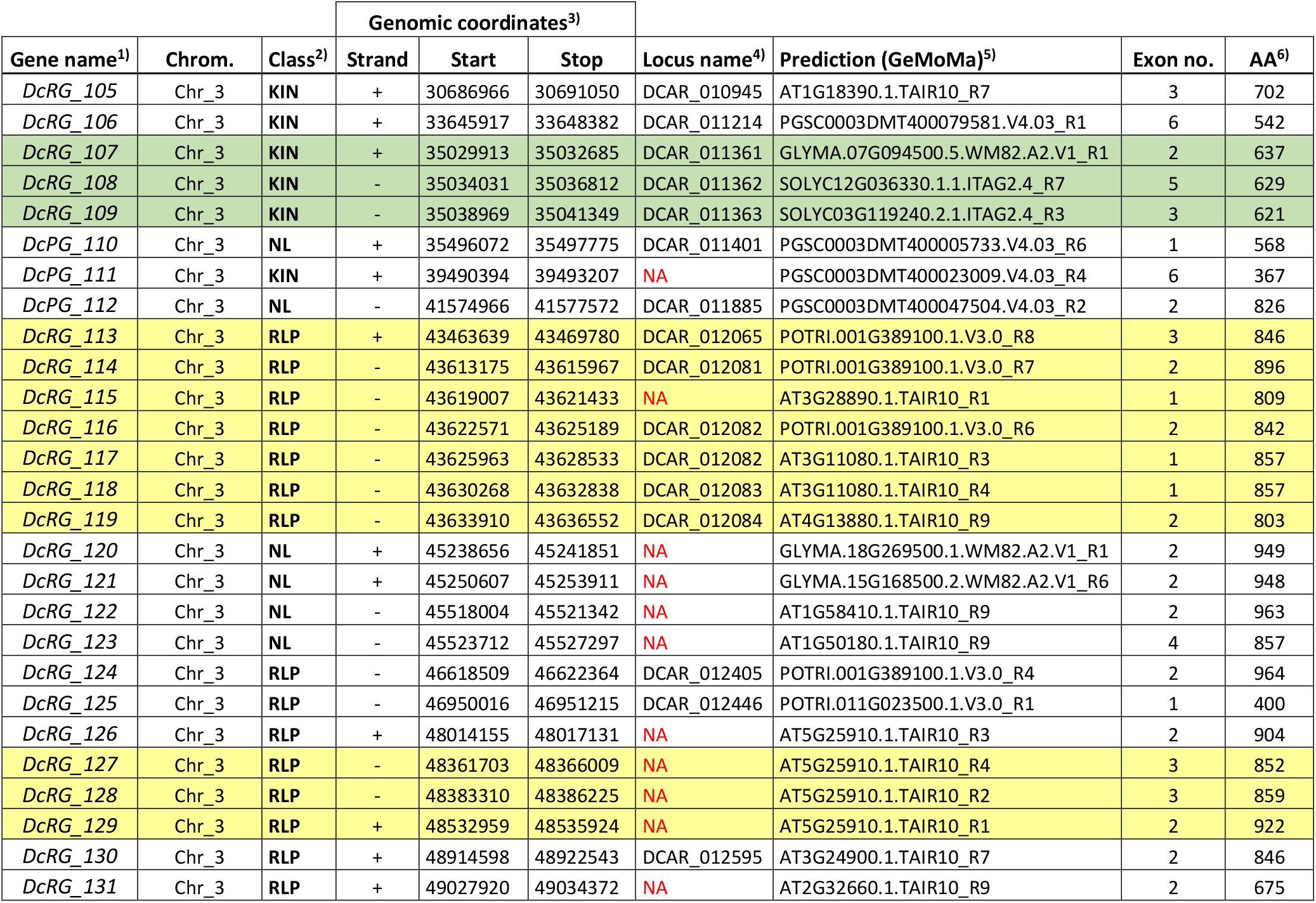
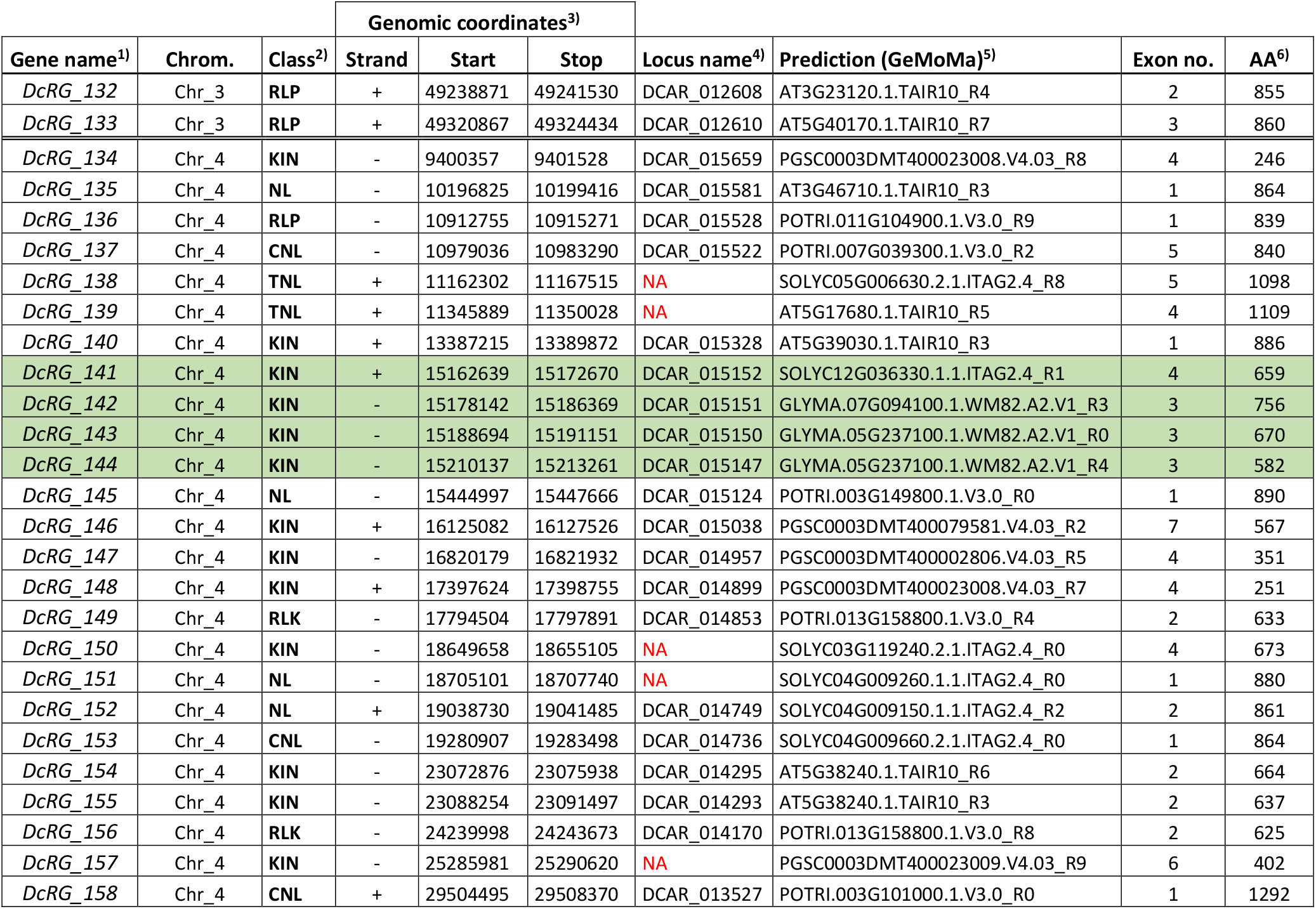
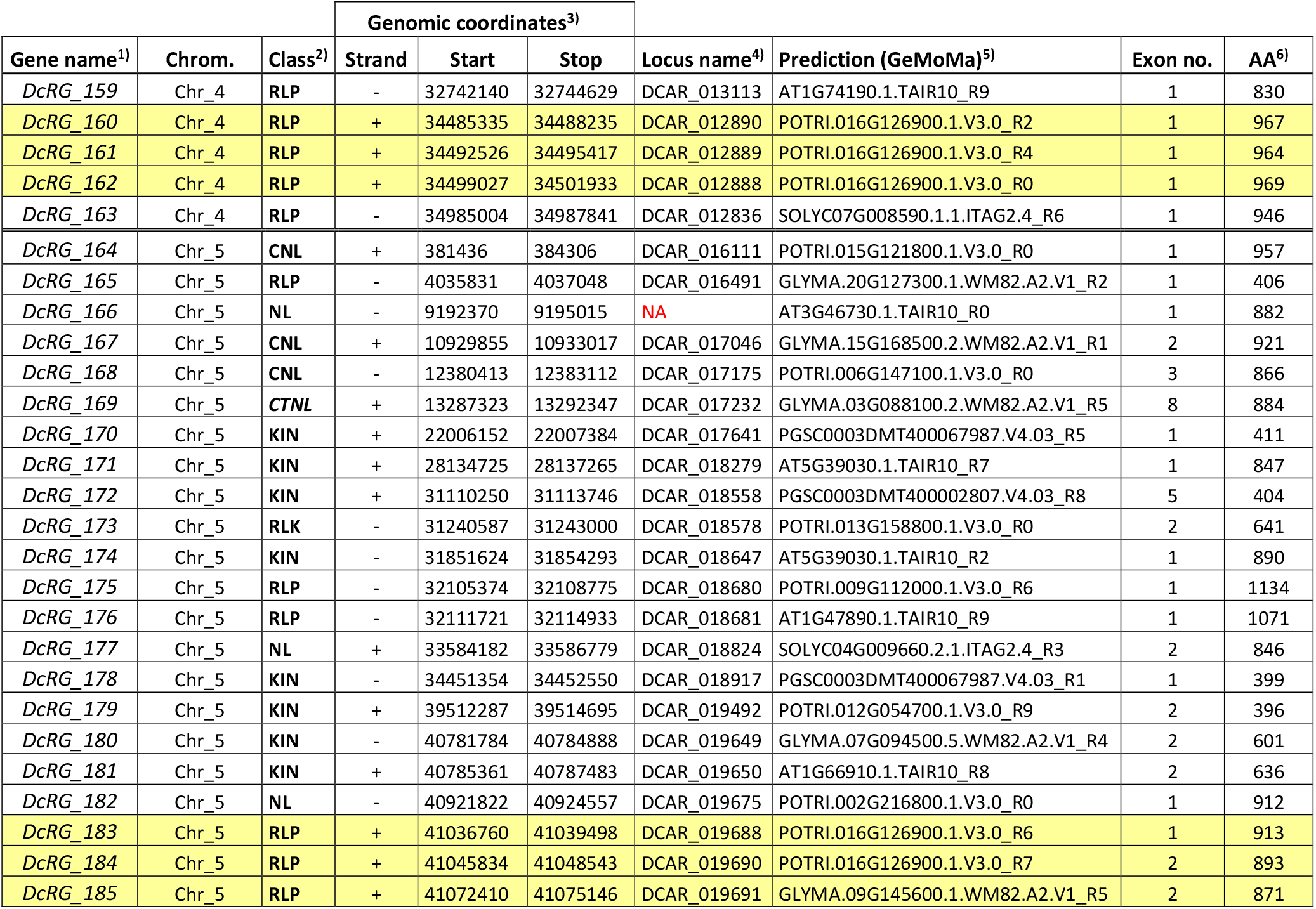
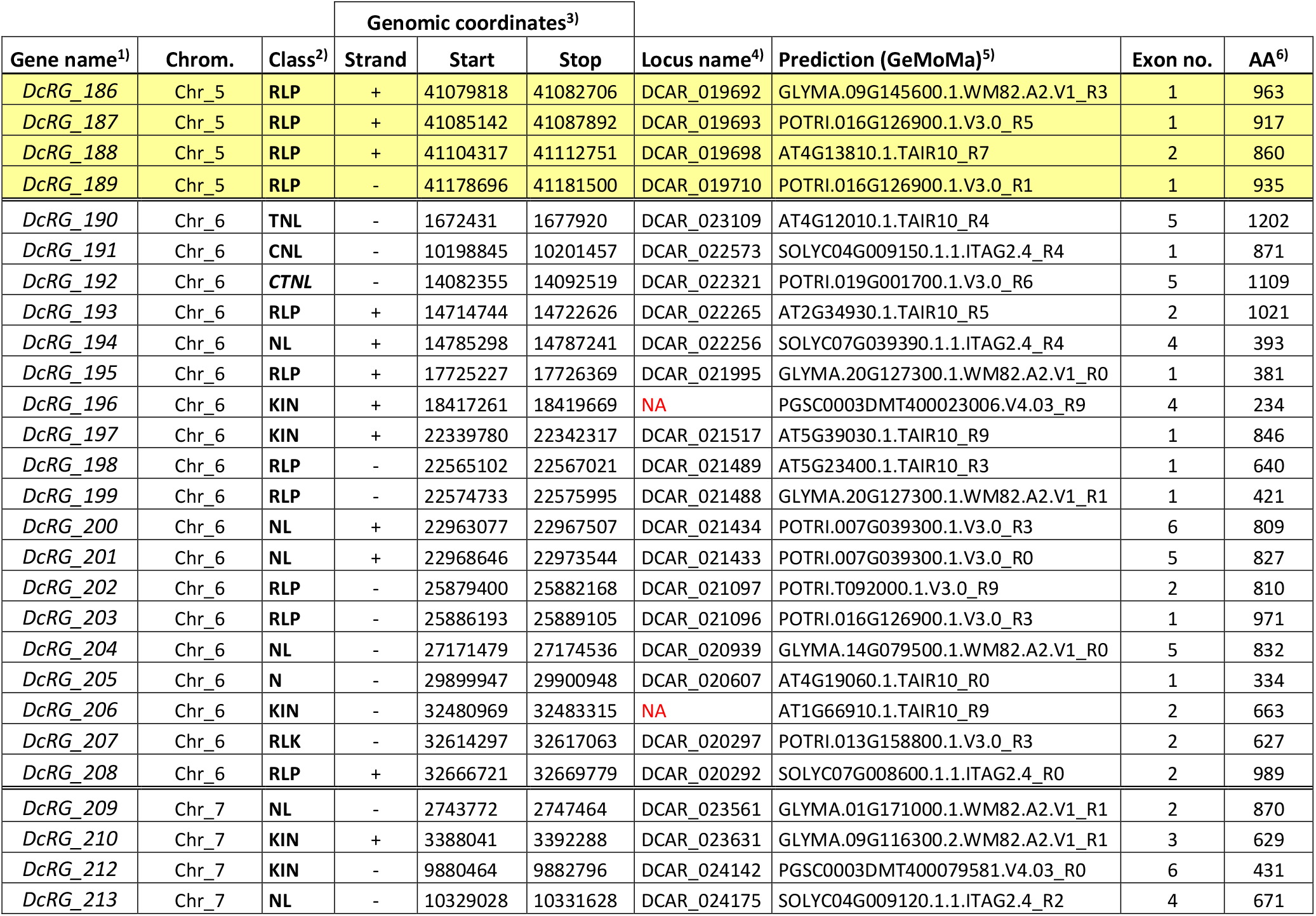
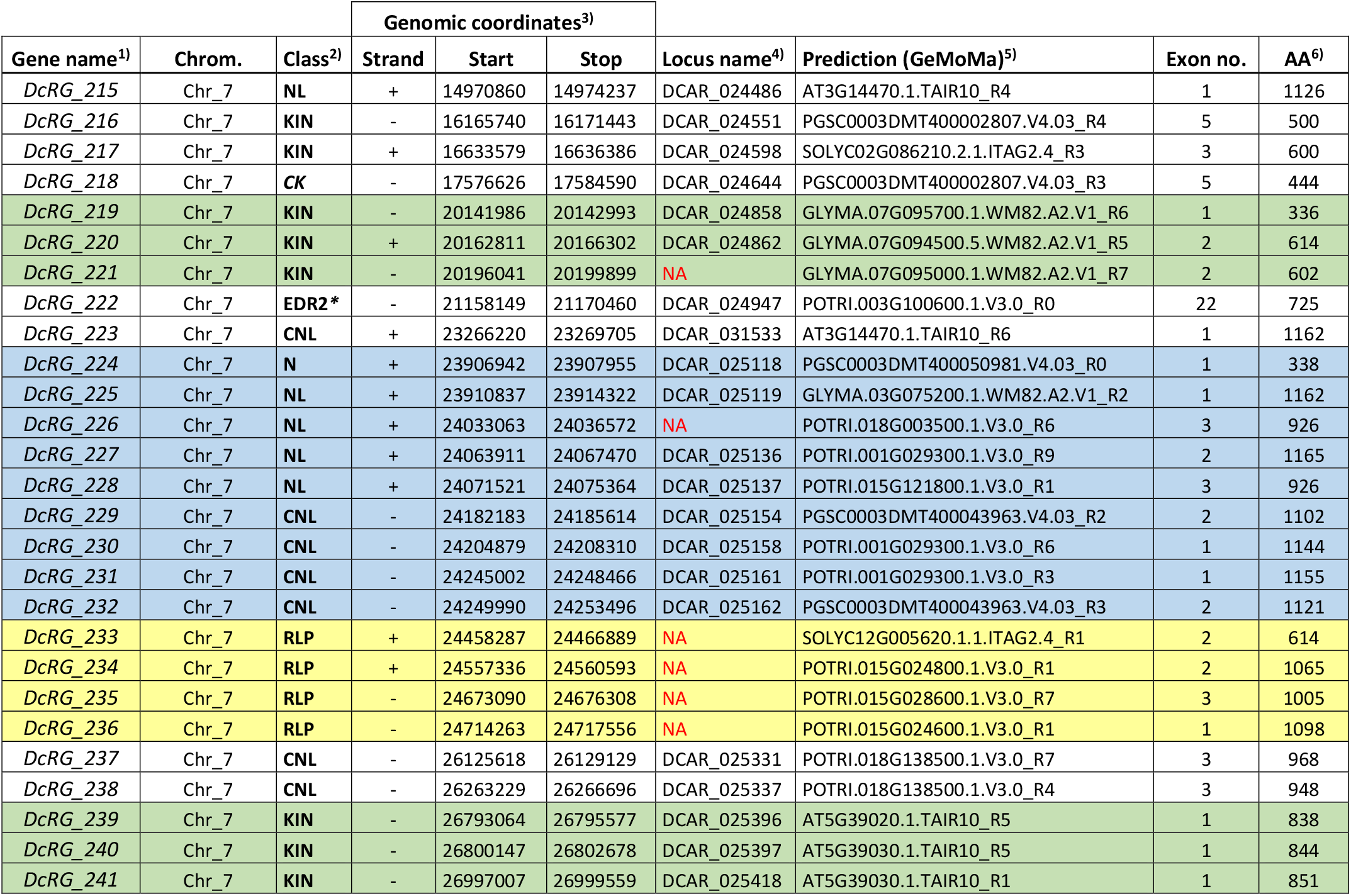
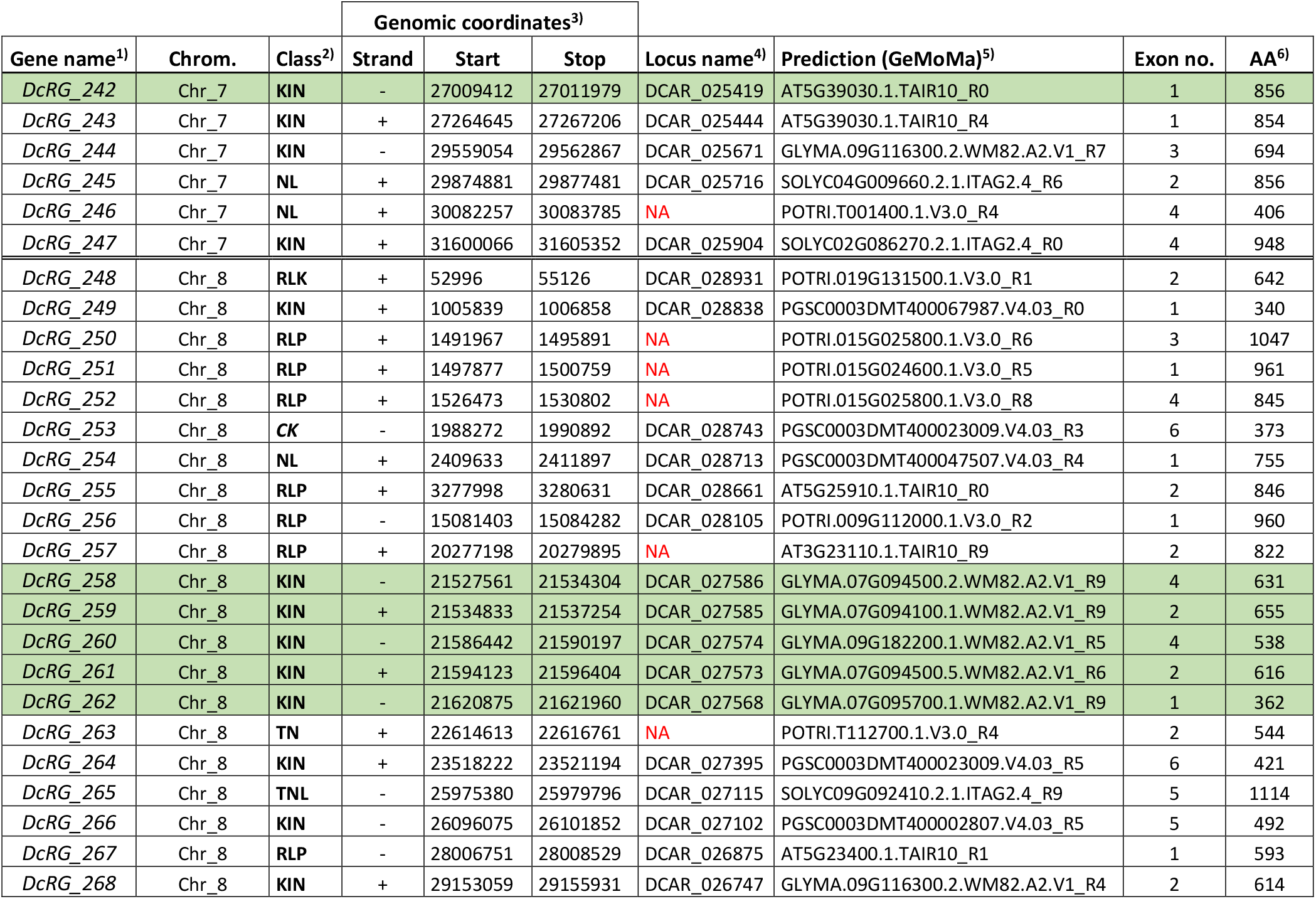
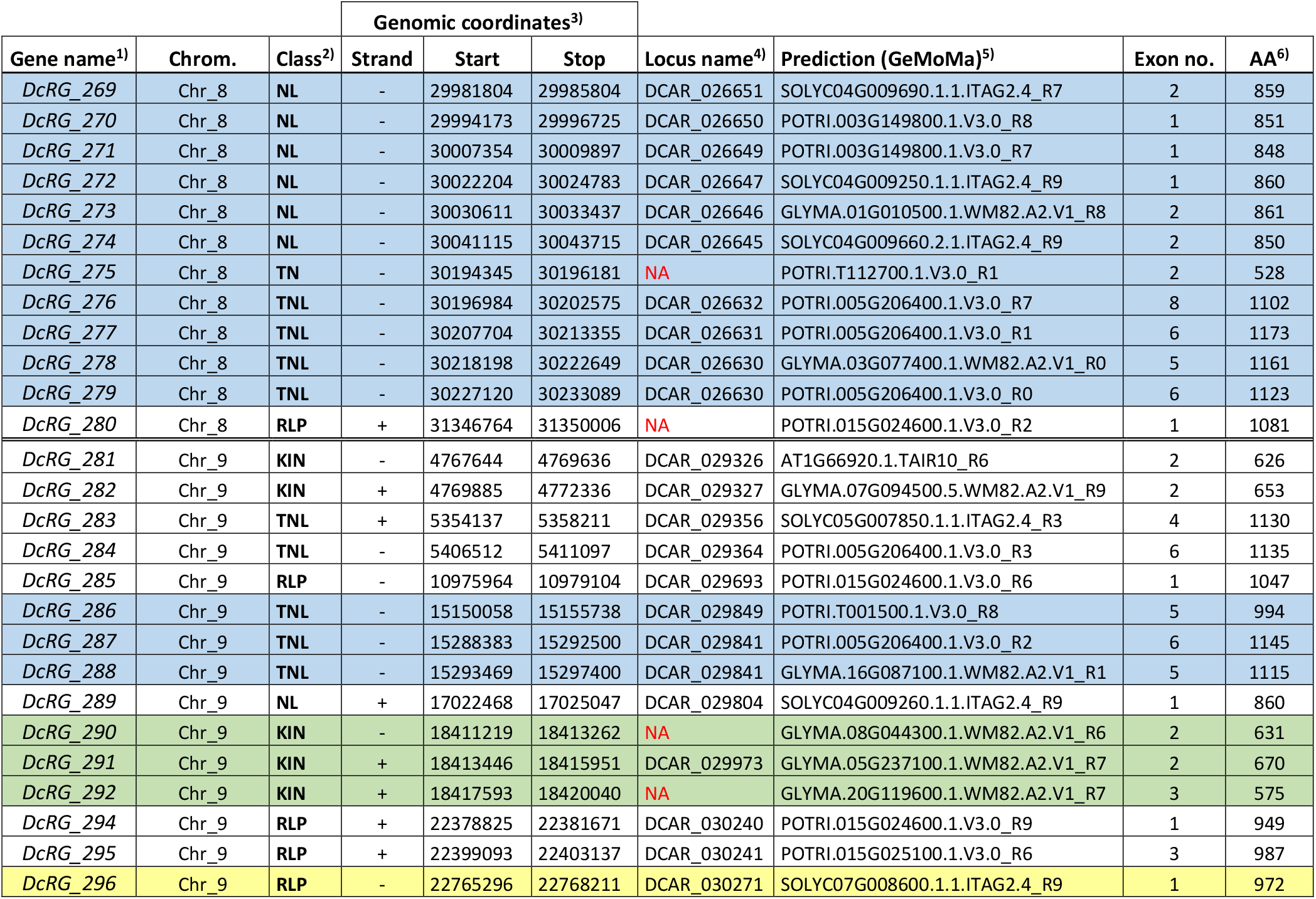
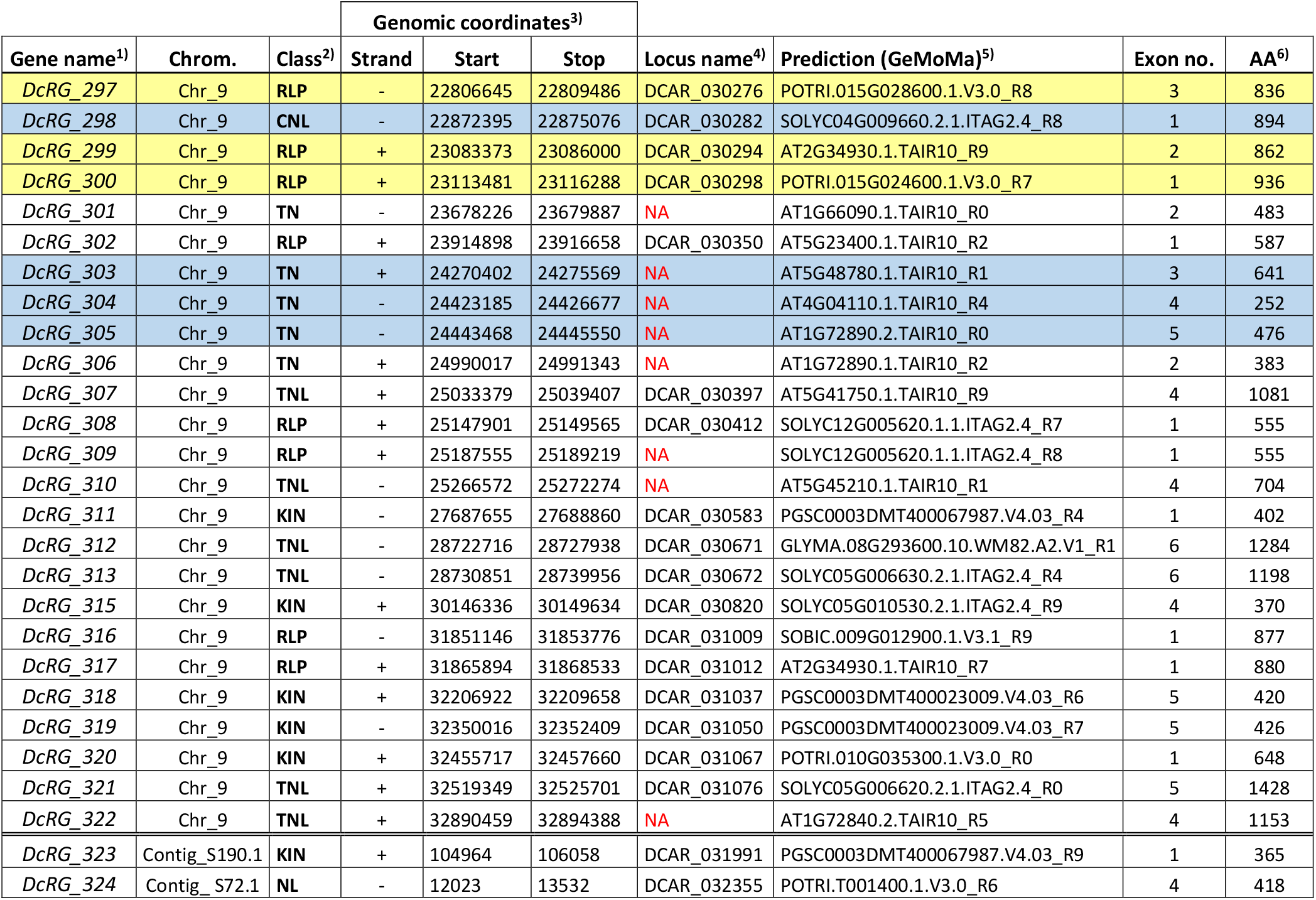
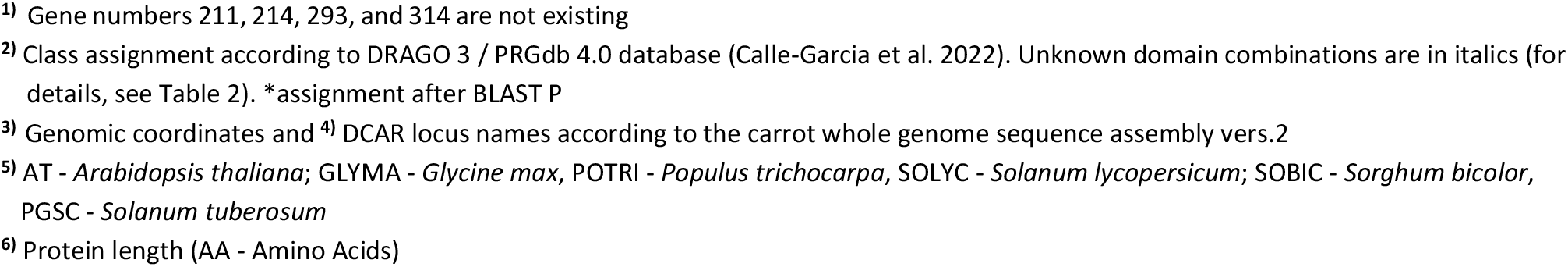
List of 320 *Daucus carota* R gene models sorted by their physical position on the assembled nine carrot chromosomes according to the carrot whole genome sequence assembly vers.2 (Iorizzo et al. 2016). NLR gene clusters are shaded in blue, RLP clusters in yellow, and KIN clusters in green, respectively (for cluster definition, see methods). NA – Not annotated in carrot genome sequence vers.2

